## Supplemental Information Tables 1-3 and Figures 1-10 for "Convergent Allostery in Ribonucleotide Reductase"

[1]Department of Chemistry and Chemical Biology, Cornell University, Ithaca, NY 14853; [2]Department of Chemistry, Princeton University, Princeton, NJ 08544; [3]Department of Chemistry, Massachusetts Institute of Technology, Cambridge, MA 02139; [4]Institute for Quantitative Biomedicine, Rutgers University, Piscataway, NJ 08854; [5]Department of Biochemistry and Molecular Biology, Oregon Health & Science University, Portland, Oregon, 97239

*Corresponding author:* Nozomi Ando, 297 Physical Sciences Building, Ithaca, NY 14853,

, (607) 255-9454

**Supplementary Tables 1-3**

**Supplementary Figures 1-10**

**Supplementary Table 1. Summary of SAXS experiments**

| **Experiment** | **Type** | **Figure** |
|---|---|---|
| 80 µM wt holo-NrdE in assay buffer with 1% glycerol | SEC | 2a |
| 40 µM wt apo-NrdE in assay buffer with 5% glycerol | SEC | 2a |
| 40 µM wt holo-NrdE + 100 µM dATP, 0.5 mM CDP in assay buffer with 1% glycerol | SEC | 2a, S2b |
| 4 µM wt holo-NrdE + 0-50 µM dATP, 1 mM CDP in assay buffer with 1% glycerol | Titration | 2b, S2a |
| 4 µM wt apo-NrdE + 0-50 µM dATP, 1 mM CDP in assay buffer with 5% glycerol | Titration | 2b, S2a |
| 4 µM C382S holo-NrdE + 0-20 µM Fe-NrdF, 50 µM dATP, and 1 mM CDP in assay buffer with 5% glycerol | Titration | 2c |
| 4 µM wt holo-NrdE + 0-1 mM dATP, 1 mM CDP in assay buffer with 1% glycerol | Titration | 3a |
| 4 µM wt apo-NrdE + 0-1 mM dATP, 1 mM CDP in assay buffer with 5% glycerol | Titration | 3a |
| 40 µM wt holo-NrdE + 1 mM ATP, 0.5 mM CDP in assay buffer with 1% glycerol | SEC | S5a |
| 4 µM wt holo-NrdE + 0-15 mM dATP, 50 µM dATP, 1 mM CDP in assay buffer with 1% glycerol | Titration | 3b |
| 4 µM wt apo-NrdE + 0-15 mM dATP, 50 µM dATP, 1 mM CDP in assay buffer with 5% glycerol | Titration | 3b |
| 4 µM wt holo-NrdE + 0-500 µM TTP, 3 mM ATP in assay buffer with 1% glycerol | Titration | 3c |
| 4 µM wt apo-NrdE + 0-500 µM TTP in assay buffer with 5% glycerol | Titration | 3c |
| 80 µM wt holo-NrdE + 250 µM TTP, 1 mM ATP in assay buffer with 1% glycerol | SEC | S5b |
| 4 µM wt holo-NrdE + 0-500 µM TTP in assay buffer with 1% glycerol | Titration | 3d |
| 40 µM wt holo-NrdE + 100 µM TTP in assay buffer with 1% glycerol | SEC | S2c |
| 80 µM C382S holo-NrdE + 80 µM Mn-NrdF, 1 mM ATP, 250 µM TTP in assay buffer with 5% glycerol | SEC | 5a, S10 |

Assay buffer = 50 mM HEPES pH 7.6, 150 mM NaCl, 15 mM $MgCl_2$, 1 mM TCEP, and 1 or 5% (w/v) glycerol. To prevent changes in nucleotide concentrations, C382S NrdE was used in all experiments where NrdF was added to the mix.

**Supplementary Table 2. Cryo-EM data collection, refinement and validation statistics**

| | #1 NrdEF<br>(EMDB-9272)<br>(PDB 6MW3) | #2 NrdE<br>(EMDB-9293)<br>(PDB 6MYX) |
|---|---|---|
| **Data collection and processing** | | |
| Magnification | 130,000× | 36,000x |
| Voltage (kV) | 200 | 200 |
| Electron exposure (e–/Å$^2$) | 5.1 to 11.7 | 20.0 |
| Defocus range (μm) | 0.8 to 2.9μm (95%) | 1.0 to 3.0 μm |
| Pixel size (Å) | 1.05 | 1.505 |
| Symmetry imposed | None | None |
| Initial particle images (no.) | 126,224 | 85,532 |
| Final particle images (no.) | 126,224 | 85,532 |
| Map resolution (Å)<br>FSC threshold | 4.65<br>(0.143) | 4.8<br>(0.143) |
| Map resolution range (Å) | 4.3-10.6 (95%) | 4.6-9.0 (95%) |
| **Refinement** | | |
| Initial model used (PDB code) | PDB 6CGL & 6MT9 | NrdEF structure |
| Model resolution (Å)<br>FSC threshold | 4.66<br>(0.5) | 5.9<br>(0.5) |
| Map sharpening *B* factor (Å$^2$) | -155 | -100 |
| Model composition<br>Non-hydrogen atoms<br>Protein residues<br>Ligands | <br>22,189<br>21,949<br>dATP | <br>55,020<br>6,730<br>dATP |
| *B* factors (Å$^2$)<br>Protein<br>Ligand | <br>122.7<br>103.1 | <br>214.01<br>161.61 |
| R.m.s. deviations<br>Bond lengths (Å)<br>Bond angles (°) | <br>0.006<br>1.072 | <br>0.005<br>0.757 |
| Validation<br>MolProbity score<br>Clashscore<br>Poor rotamers (%) | <br>1.73<br>4.85<br>0.00 | <br>2.21<br>9.6<br>3.06 |
| Ramachandran plot<br>Favored (%)<br>Allowed (%)<br>Disallowed (%) | <br>92.20<br>7.80<br>0.00 | <br>95.13<br>4.87<br>0.00 |

**Supplementary Table 3**. **Diffraction data and model refinement statistics for *B. subtilis* NrdE protein crystals.**

| **Data Collection**[a] | Disulfide-trapped NrdE[b] (mainly oxidized) (PDB 6MT9) | X-ray-reduced NrdE[b] (partially reduced) (PDB 6MVE) | NrdE with empty M-site[c] (PDB 6MV9) |
|---|---|---|---|
| Space group | $P4_32_12$ | $P4_32_12$ | $P2_12_12_1$ |
| Unit cell (Å) | a = b = 126.34, c = 125.44 α = β = γ = 90.0 | a = b = 126.41, c = 125.49 α = β = γ = 90.0 | a = 120.26; b = 126.40, c = 128.31 α = β = γ = 90.0 |
| Wavelength (Å) | 0.9775 | 0.9775 | 0.9775 |
| Resolution range (Å) | 16.00 – 2.50 (2.60 – 2.50) | 15.99-2.55 (2.66-2.55) | 19.98-2.95 (3.07-2.95) |
| Total observations | 601897 (69514) | 439944 (54618) | 149881 (17349) |
| Total unique observations | 35371 (3958) | 33640 (4072) | 41464 (4649) |
| $R_{merge}$ | 0.175 (1.445) | 0.162 (1.500) | 0.180 (0.763) |
| $R_{pim}$ | 0.041 (0.333) | 0.046 (0.424) | 0.108 (0.452) |
| ⟨I/σ(I)⟩ | 14.7 (2.2) | 11.8 (1.9) | 5.4 (1.6) |
| $CC_{1/2}$ | 0.998 (0.734) | 0.998 (0.745) | 0.978 (0.441) |
| Completeness (%) | 99.2 (99.9) | 99.6 (100.0) | 99.2 (99.8) |
| Multiplicity | 17.0 (17.6) | 13.1 (13.4) | 3.6 (3.7) |
| **Refinement Statistics** | | | |
| Resolution range (Å) | 2.50-16.00 | 2.55-15.99 | 2.95-19.98 |
| Reflections (total) | 35313 | 33577 | 41404 |
| Reflections (test) | 2778 | 2642 | 950 |
| Total atoms refined | 5690 | 5621 | 10896 |
| $R_{work}$ / $R_{free}$ | 0.178/0.218 | 0.175/0.218 | 0.207/0.239 |
| RMSD of bond lengths (Å)/ angles (°) | 0.006/0.919 | 0.008/1.070 | 0.003/0.619 |
| Ramachandran plot favored/allowed (%) | 97.9/2.1 | 97.3/2.7 | 96.1/3.9 |
| Mean B value for all atoms ($Å^2$) | 48.0 | 54.0 | 50.0 |

[a] Data collection values in parentheses refer to the high-resolution shell; [b] Crystal was grown with GDP substrate in the crystallization solution; [c] Crystal was grown with CDP substrate in the crystallization solution.

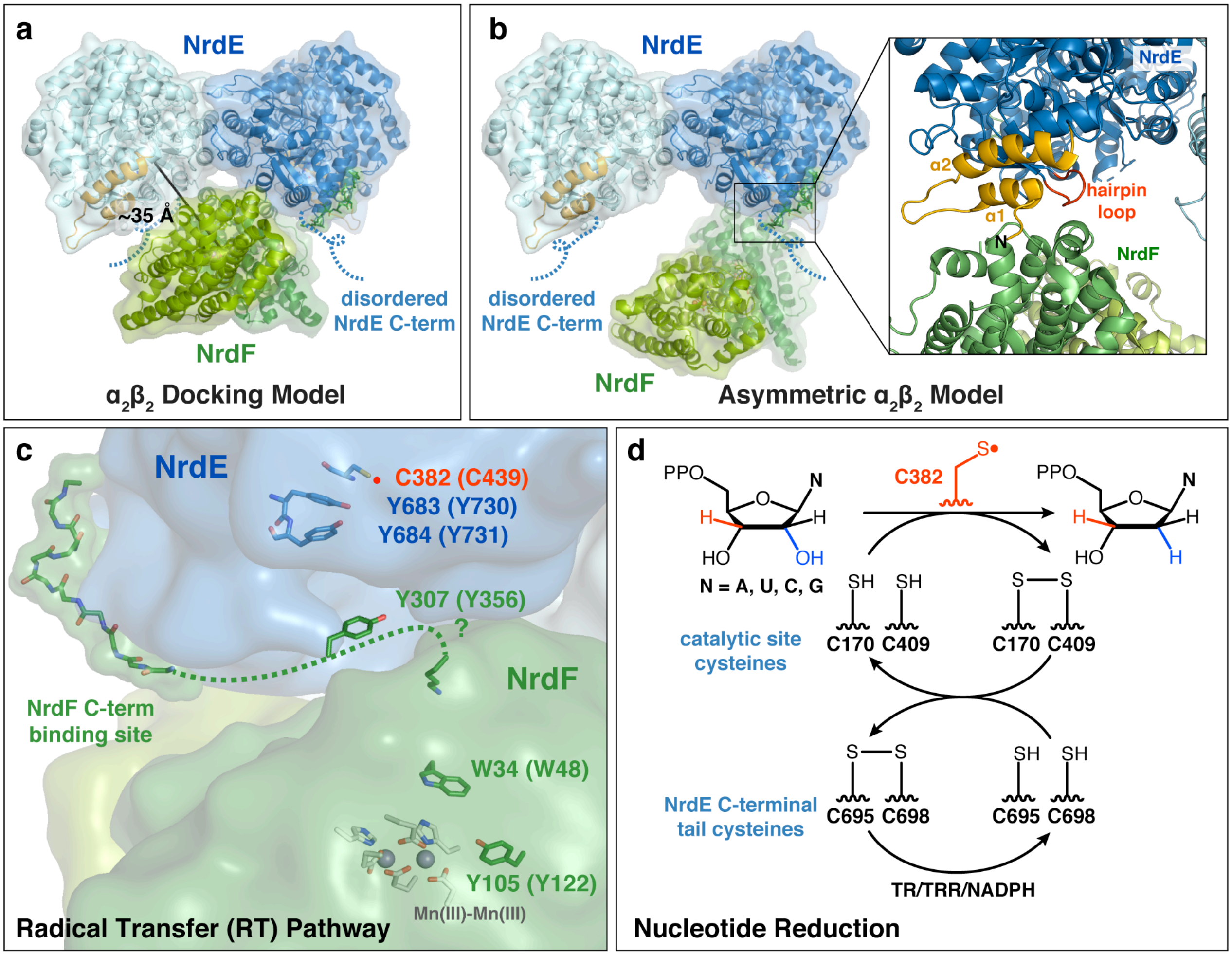

**Supplementary Figure 1. Current structural and mechanistic models of class I RNRs and the roles of the C-termini. (a)** In class I RNRs, a compact $\alpha_2\beta_2$ configuration is thought to form for radical transfer (RT). Biochemical and low-resolution structural studies on class Ia RNRs support this docking model[16,18,28,34]. **(b)** A crystal structure of the *S. typhimurium* class Ib RNR depicts NrdF bound asymmetrically to a single monomer of the NrdE S-dimer (PDB: 2BQ1)[33], forming an interface at the partial ATP-cone (orange cartoon in inset) and a β-hairpin loop (red cartoon in inset). Although this conformation would not be capable of RT, it was proposed to represent a conformation needed for the re-reduction of the active site by the disordered α C-terminus (blue dotted line). **(c)** For each turnover, the catalytic thiyl radical (C382, red) is generated by reversible long-range proton-coupled electron transfer (PCET) over a specific pathway[16]: Y105•/[W34]/Y307 in β to Y684/Y683/C382 in α (*E. coli* Ia numbering shown in parentheses). Brackets indicate that the direct involvement of W34 has not yet been demonstrated. The β C-terminus binds α for complex formation. This tail also contains Y307, the residue that is responsible for RT across the subunit interface, but the residue's location is unknown. This region of the β C-terminus is disordered in all RNR structures (dotted green line). Here, the RT pathway is oriented according to the docking model in (a). **(d)** Once the cysteine radical (red) is generated, nucleotide reduction proceeds using two additional catalytic-site cysteines as reducing equivalents. Two cysteines on the α C-terminus re-reduce the catalytic site and are ultimately reduced by thioredoxin (TR), thioredoxin reductase (TRR), and NADPH.

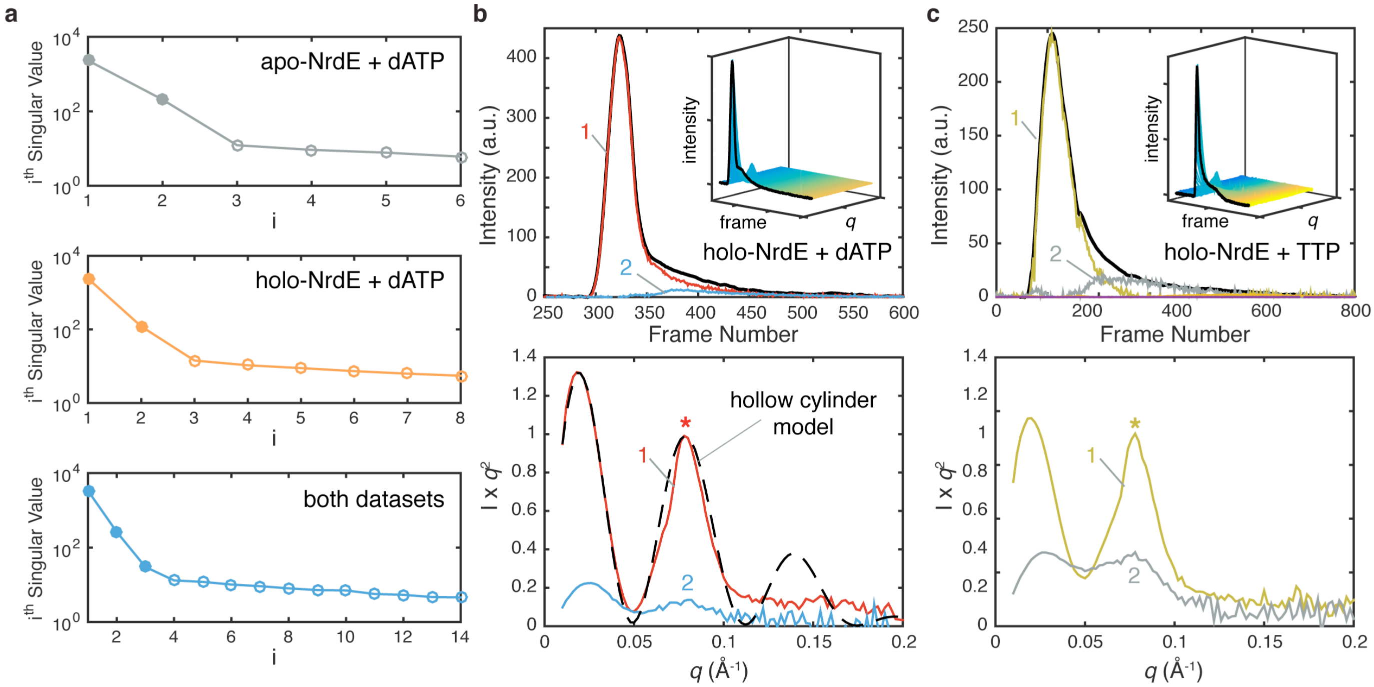

**Supplementary Figure 2. SAXS analyses of dATP- and TTP-induced formation of an extended NrdE oligomer. (a)** Singular value decomposition (SVD) was performed on dATP titrations shown in Fig. 2b for apo-NrdE (*top*), holo-NrdE (*middle*), and both datasets combined (*bottom*). Significant singular values are shown as closed circles. Apo- and holo-NrdE each have two significant singular values, whereas the combined dataset has three, indicating that the final state of the dATP titration is the same for both samples. **(b)** SEC-SAXS was performed on holo-NrdE with 100 µM dATP (*top, inset*). Evolving factor analysis (EFA) of the dataset (overall elution shown as black curve) yielded two scattering components with the major species eluting first (*top*, red) and having scattering features (*bottom*, red) that can be described by a hollow cylinder model with inner radius of 35.2 Å, outer radius of 62 Å, and length of 950 Å (black dashed line). SAXS profiles are shown in Kratky representation to emphasize mid-*q* features, such as the prominent second peak (star). **(c)** SEC-SAXS was performed on holo-NrdE with 100 µM TTP in the absence of ATP (*top, inset*), and 4 sequentially eluting species were separated using EFA. The major component elutes first (*top*, yellow) and has a similar shape to the dATP-induced oligomer, including a prominent second peak (*bottom*, star). Components 3 and 4 are not visible in the elution as they are minor contributions. The secondary peak of the extended NrdE oligomer is visible even in the raw SEC-SAXS data (*insets* of (b) and (c)).

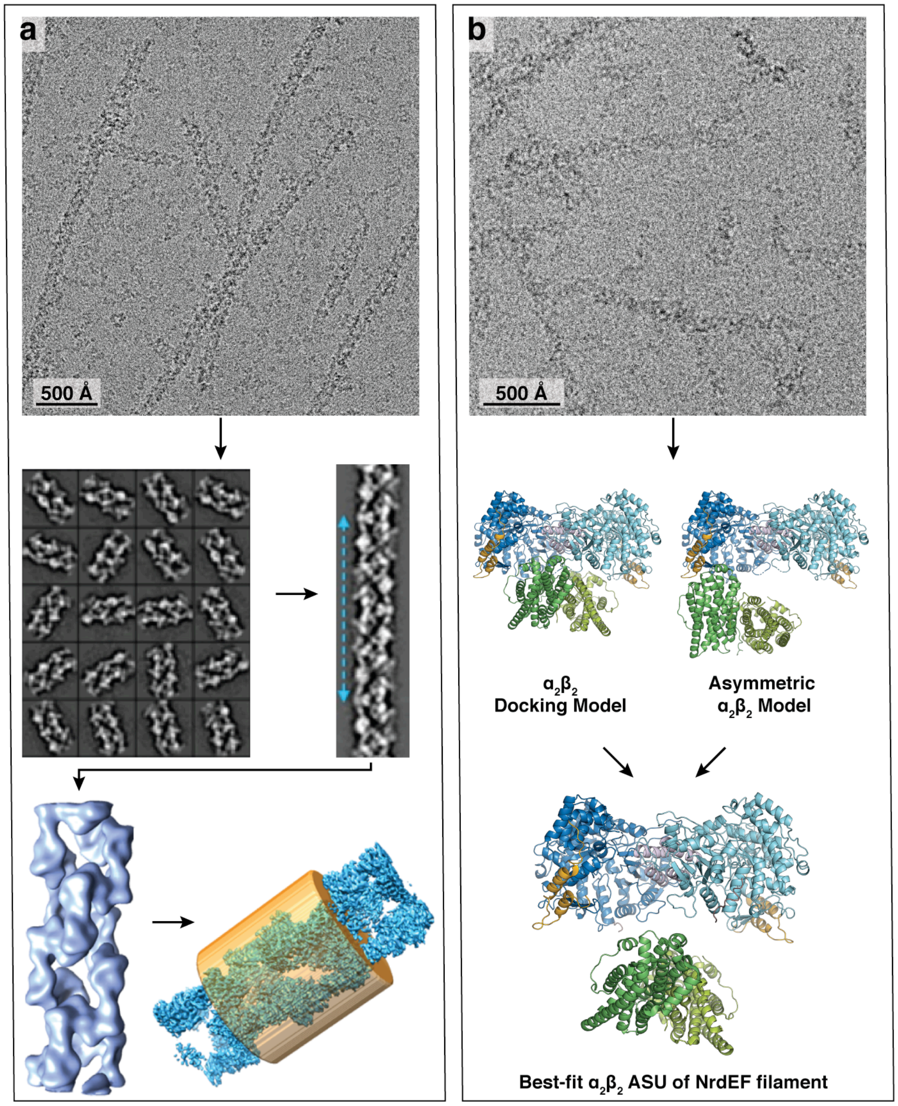

**Supplementary Figure 3. Cryo-EM analyses of the dATP-induced NrdE and NrdEF filaments. (a)** Single-particle reconstruction and refinement of the 4.8-Å NrdE filament map. A representative cryo-EM micrograph is shown at the top. Class averages of picked particles were used to build a full helical segment (*middle*). Initial helical reconstruction converged to a 5.8-Å structure that was further refined to 4.8 Å via masking and local refinement (*bottom*). **(b)** *Ab initio* single-particle reconstruction of the NrdEF filament led to a model with a large gap between the NrdE and NrdF subunits. To test this model, the map was re-refined using two starting models: a symmetric $\alpha_2\beta_2$ based on an *E. coli* docking model[18] and an asymmetric $\alpha_2\beta_2$ based on an *S. typhimurium* crystal structure (PDB: 2BQ1)[33]. In both cases, refinement converged on a map in which there was a large gap between aligned NrdE and NrdF dimer structures. A representative cryo-EM micrograph is shown at the top.

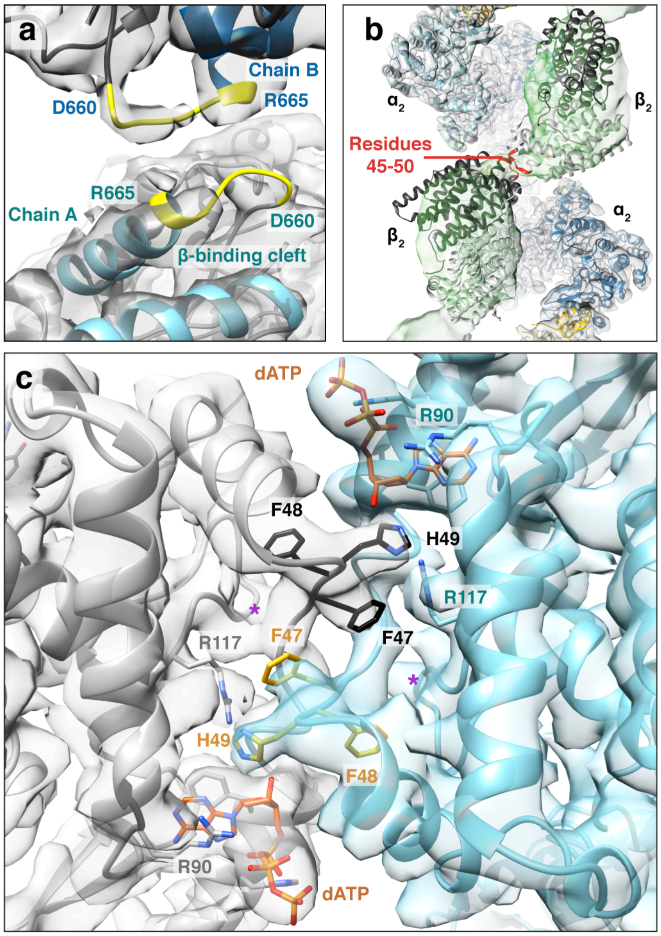

**Supplementary Figure 4. Interfaces observed in cryo-EM structures of dATP-induced filaments. (a)** The NrdE filament forms the double-helix interface (yellow cartoon) at the β-tail-binding cleft (blue cartoon), which helps explain dissociation of the double-helix upon addition of NrdF. EM density is shown in grey (threshold = 4.24). **(b)** Rigid-body fitting of $\beta_2$ (black and white) into non-NrdE density (green, threshold = 0.45) in the 4.7-Å NrdEF map. The orientation with the highest map-model correlation leads to a symmetric interface between adjacent NrdF dimers via residues 45-50 (red). $\beta_2$ was modeled using a crystal structure of *B. subtilis* NrdF (PDB: 4DR0)[21]. **(c)** The I-dimer interface is formed by interlocking loops (residues 45-50, gold and black) from both chains. At the center of this loop, side-chain density for F47 and H49 is observed extending across the interface. Individual chain density of the NrdEF map is shown here in grey and cyan (threshold = 1.44). The M-site (purple star) is empty.

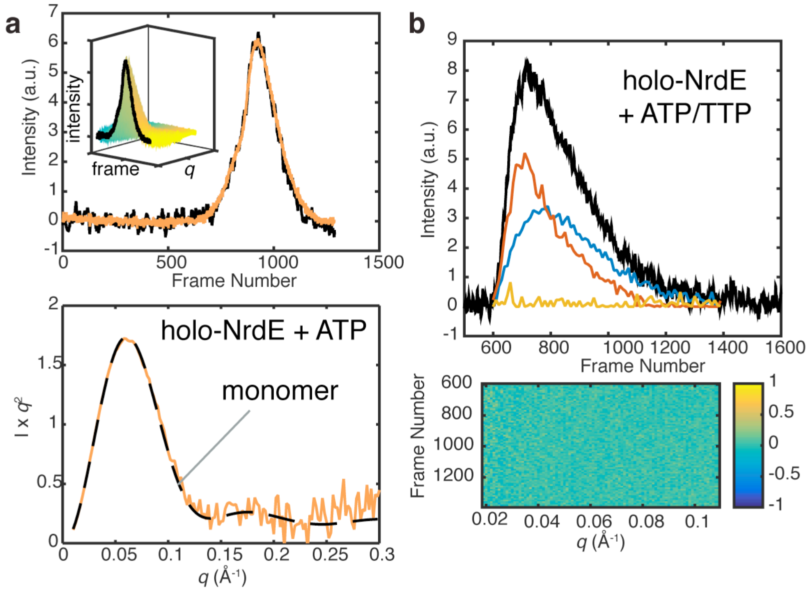

**Supplementary Figure 5. SAXS experiments reveal that ATP disrupts the I-dimer interface and specificity effectors induce S-dimer formation. (a)** SEC-SAXS was performed on holo-NrdE with 1 mM ATP (*top, inset*). EFA of the dataset (overall elution shown as black curve) yielded a single scattering component (*top*, orange) with scattering features (*bottom*, orange) that are well described by the crystal structure of monomeric NrdE (black dash, PDB: 6CGM)[31]. **(b)** SEC-SAXS of NrdE was performed under conditions with holo-NrdE + 1 mM ATP and 250 μM TTP. The full dataset was fit to a linear sum of the theoretical scattering of the I-dimer (PDB: 6CGL)[31], S-dimer (this work, PDB: 6MT9), and monomer (PDB: 6CGM) using the program OLIGOMER[52,58]. The top panel shows the scattering contribution of S-dimer (red), I-dimer (yellow), and monomer (blue) overlaid with the low-*q* intensity (black) for the SEC elution. NrdE was found to be best explained as a mixture of S-dimer and monomer with no I-dimer component. The co-elution of these species indicates that they are able to rapidly exchange. The bottom panel shows a map of the residuals of the OLIGOMER fits as a function of frame number and *q*. The residuals are largely zero and have no *q*- or frame-dependence.

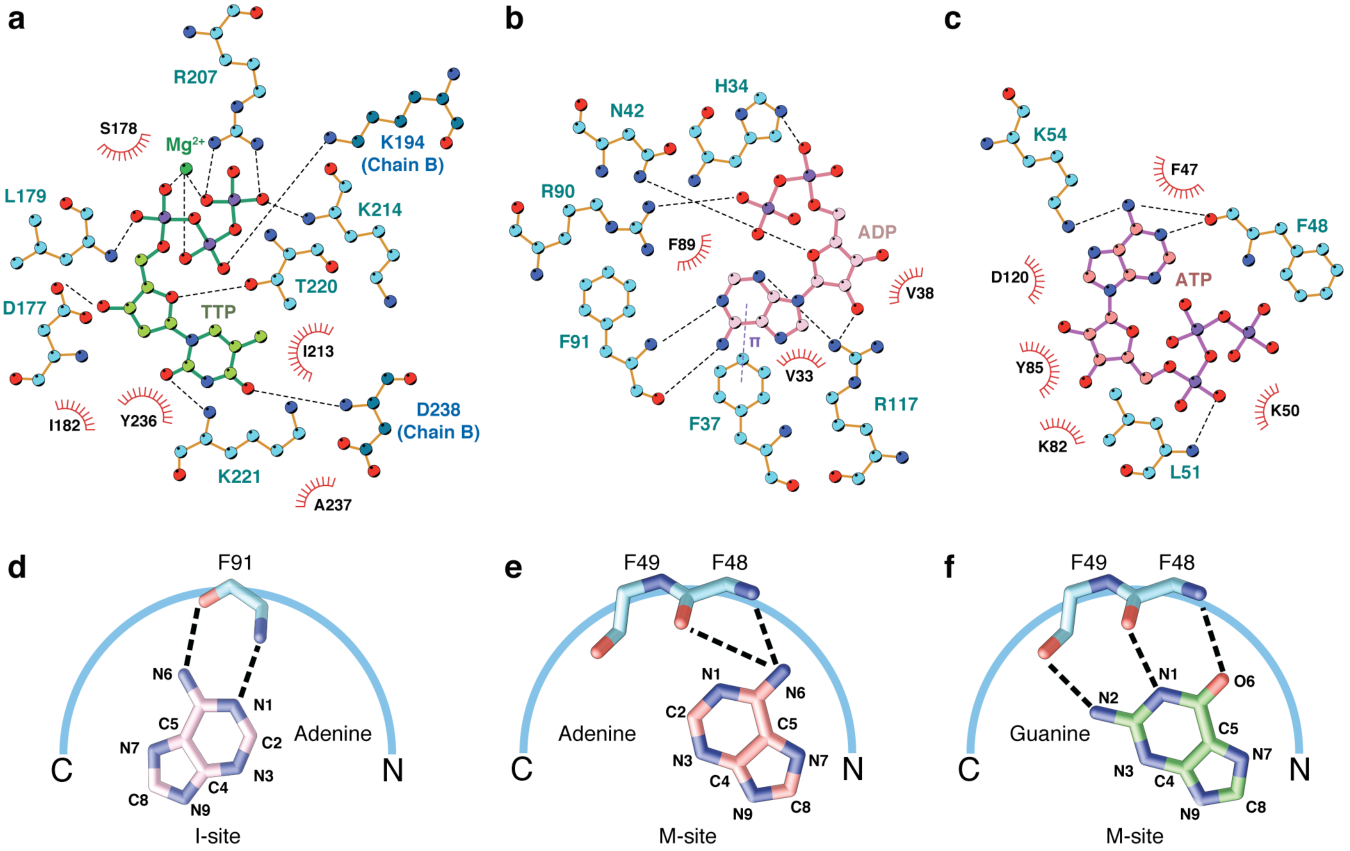

**Supplementary Figure 6. Protein-nucleotide interaction diagrams of crystal structures.** *Top row:* Ligand-interaction diagrams for our 2.50-Å NrdE S-dimer structure. Polar interactions are shown as dotted lines and hydrophobic contacts are shown as red arcs with radiating spokes. **(a)** Residues from both chains of the S-dimer interact with TTP (carbon atoms shown in green) at the S-site. **(b)** ADP (carbon atoms shown in pink) bound to the I-site. **(c)** ATP (carbon atoms shown in salmon) bound to the M-site. Unless otherwise noted, carbons, oxygens, nitrogens, and $Mg^{2+}$ are shown in light blue, red, dark blue, and dark green, respectively. *Bottom row:* Backbone-nucleobase interactions at the I- and M-sites. **(d)** In all three of our structures as well as structures with dAMP bound[31], the adenine ring of (d)AxP binds the I-site via a reverse adenine-binding interaction using the backbone of residue F91. Here, "reverse" refers to the motif having a C- to N-terminus directionality[77]. **(e)** In our 2.50- and 2.55-Å structures, F48 of the M-site forms a backbone interaction with the adenine ring of ATP, but the positioning prevents interaction of the N1 group with the M-site, as N1 cannot act as an H-bond donor to the backbone carbonyl oxygen of F48. **(f)** Although it is not supported by the electron density in our structures, modeling a guanosine nucleotide in the M-site in the same orientation as ATP reveals three favorable binding interactions. Notably, the N1 group could form an H-bond with the backbone carbonyl of F48.

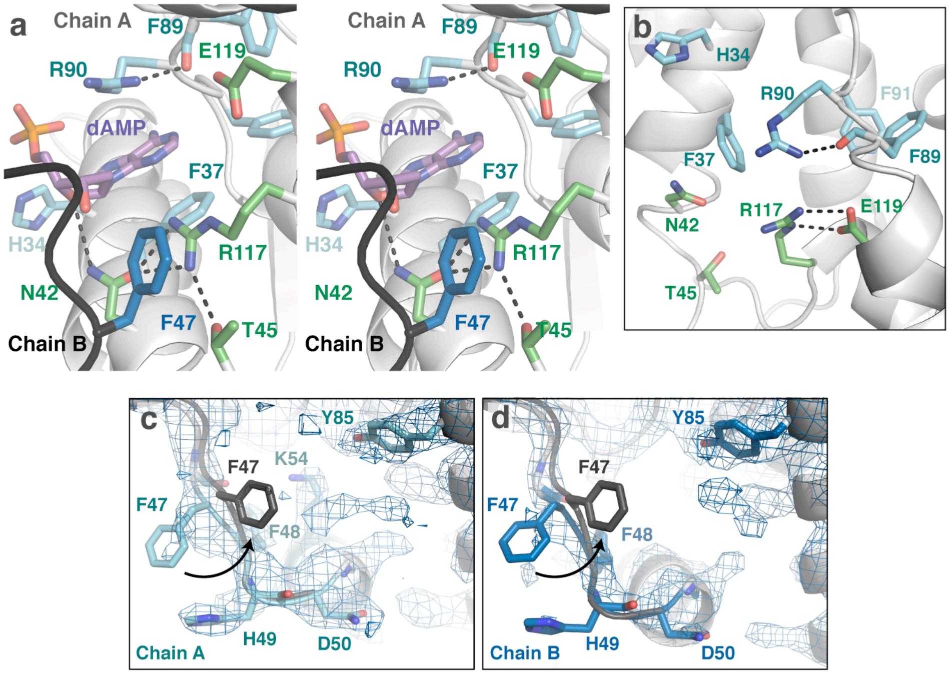

**Supplementary Figure 7. The roles of F47 at the I- and M-sites. (a)** A stereoview of the I-site in the crystal structure of the dAMP-bound NrdE I-dimer (PDB: 6CGL)[31] reveals that the F47 phenyl ring of one chain (B) is positioned to form a π-cation interaction with R117 of the opposing chain (A). This interaction allows the F47-loop from opposing chains to interlock and form an I-dimer interface. **(b)** In the apo-NrdE structure (PDB: 6CGM)[31], R117 interacts with E119 rather than N42 and T45, thus mimicking the ribonucleotide-bound conformation as seen in Figure 4e. (**c-d**) $2F_o$-$F_c$ electron density map for our 2.95-Å NrdE S-dimer structure obtained with CDP in the crystallization condition is shown as a blue mesh contoured at 1σ. No convincing electron density for a nucleotide is found adjacent to the F47-loop in either (**c**) chain A (cyan) or (**d**) chain B (blue). Consistent with having empty M-sites, the F47-loop is slightly disordered, particularly in chain B, but there is a clear difference in both the side-chain and backbone positions when compared to the F47-loop of our 2.50-Å structure (black cartoon overlay). In our 2.50-Å structure, ATP is bound in the M-site, causing F47 to swing inward (arrow). This is expected to disfavor I-dimer formation.

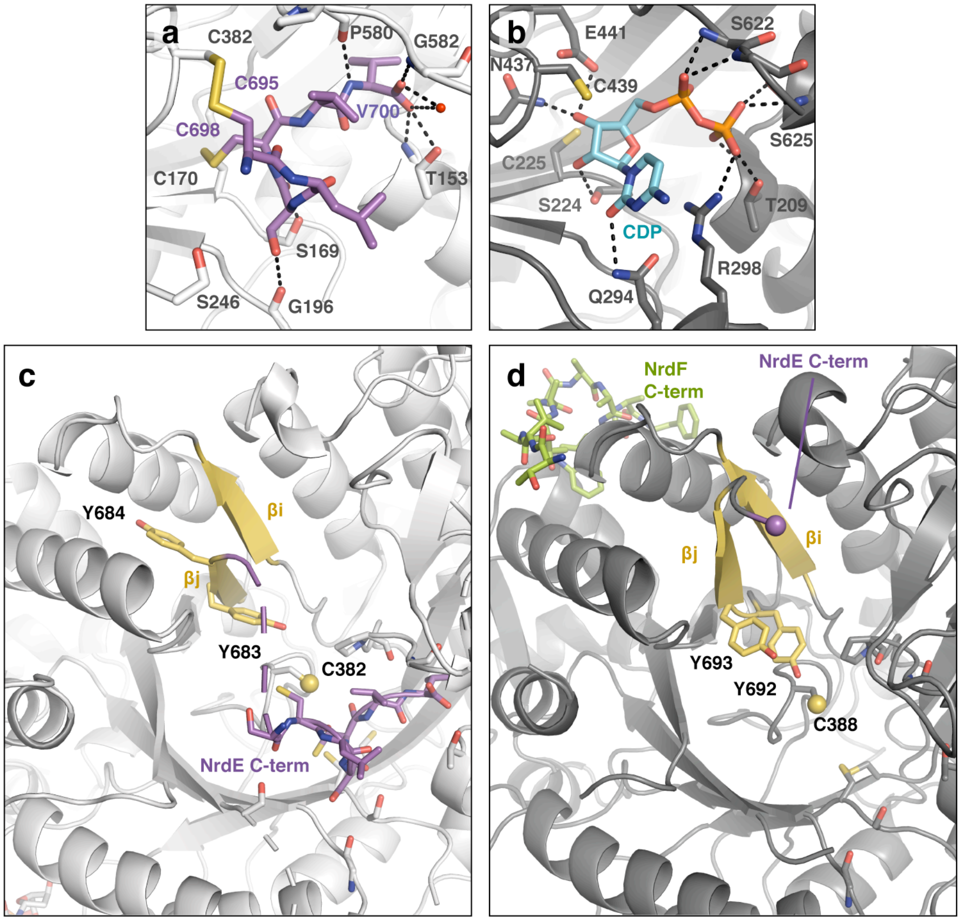

**Supplementary Figure 8. Crystal structures provide insight into control of radical transfer, nucleotide reduction, and re-reduction. (a)** The catalytic site with the oxidized NrdE C-terminus in the 2.50-Å S-dimer structure (PDB: 6MT9). The C-terminus binds in the same location as a nucleotide substrate and has several similar interactions. Notably, the terminal carboxylate of the protein interacts with charged and polar residues in a manner reminiscent of the β-phosphate of the substrate. **(b)** The crystal structure of *E. coli* class Ia RNR shown in the same orientation reveals polar interactions observed between the catalytic site residues and the substrate, CDP (PDB: 5CNS)[19]. **(c)** In our structures of the NrdE catalytic site, the β-strand containing the tyrosine dyad (Y683/Y684) is partially unzipped from the β-sheet (yellow) to allow for the NrdE C-terminus (purple sticks) to reach the catalytic site (disordered region shown as purple dotted line). This change in secondary structure is accompanied by the unstacking of Y684 from Y683. **(d)** In the structure of the *S. typhimurium* NrdEF complex (PDB: 2BQ1)[33], the NrdE tyrosines involved in PCET (Y692/Y693) form a stacked dyad over the catalytic cysteine (C388). The tyrosine dyad is located in the β-strand (yellow) directly preceding the NrdE C-terminal tail (purple sphere). The C-terminus of NrdF (green) is observed bound within a hydrophobic cleft on the surface of the catalytic barrel in this structure.

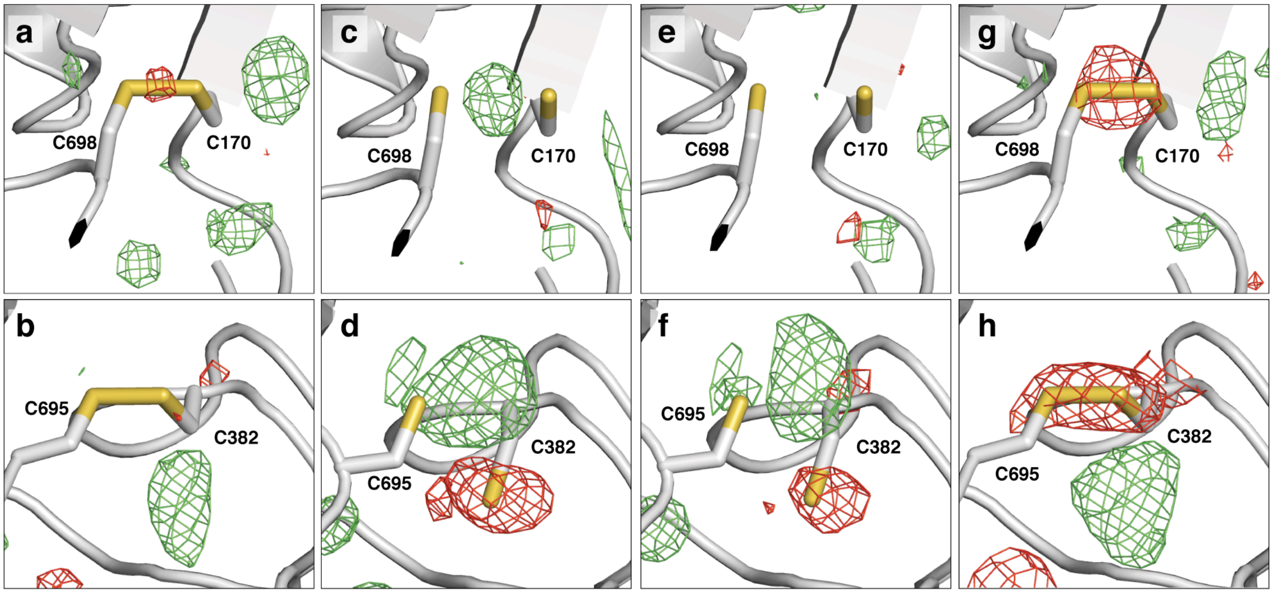

**Supplementary Figure 9. Active-site redox state of α C-terminus trapped in crystal structures of *B. subtilis* NrdE.** Both datasets show signs of partial oxidation and reduction. However, the 2.50-Å dataset is better described by an oxidized model, and the 2.55-Å dataset is better described by the reduced model. **(a-b)** $mF_o$-$DF_c$ map synthesized using the refined oxidized model and 2.50-Å dataset. **(c-d)** $mF_o$-$DF_c$ map synthesized using the refined reduced model and 2.50-Å dataset. **(e-f)** $mF_o$-$DF_c$ map synthesized using the refined reduced model and 2.55-Å dataset. **(g-h)** $mF_o$-$DF_c$ map synthesized using the refined oxidized model and 2.55-Å dataset. All maps are contoured at 3σ, with the green mesh representing positive $F_o$ density and the red negative $F_c$ density.

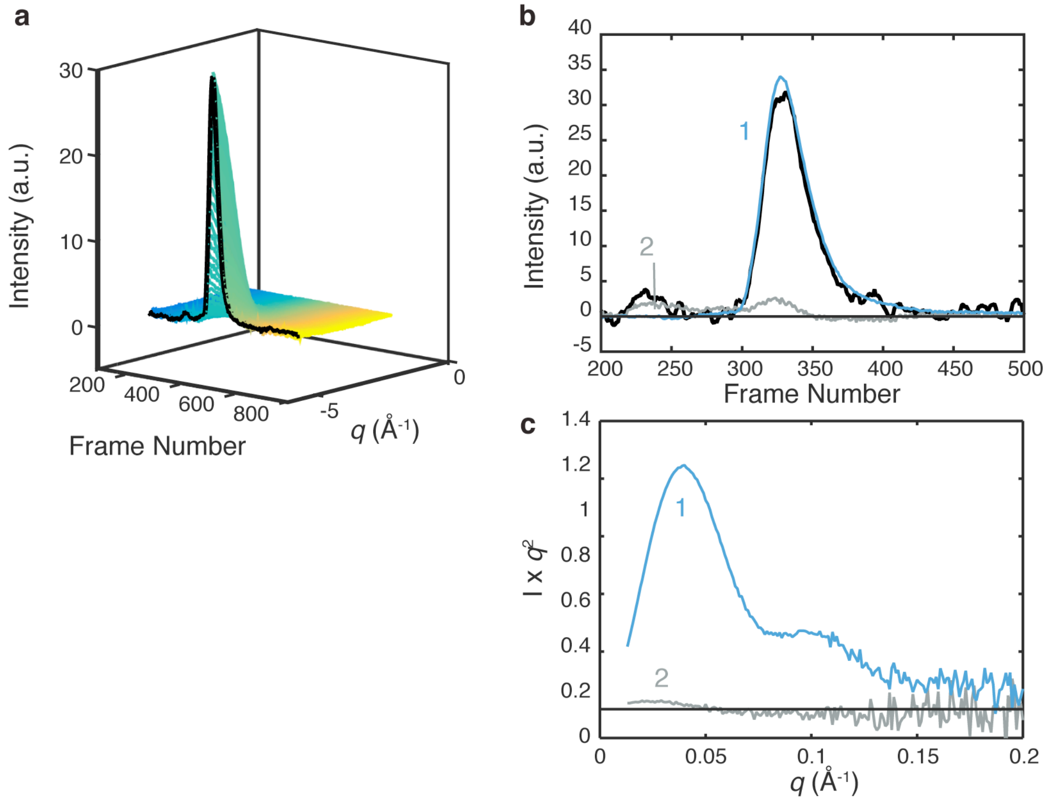

**Supplementary Figure 10. NrdEF is predominantly $\alpha_2\beta_2$ under activating nucleotide conditions. (a)** SEC-SAXS was performed on an equimolar solution of C382S holo-NrdE and Mn-NrdF pre-incubated with 1 mM ATP, 250 µM TTP and eluted under identical buffer conditions. **(b)** EFA of the overall elution profile (black) reveals one major protein component (1, blue) and a minor second component (2, grey). No additional protein components were observed eluting after the major component, suggesting that the two subunits bind strongly. **(c)** The Kratky plots of the two components separated by EFA. The minor component has a partially negative profile that does not resemble normal protein scattering and instead may represent the changing background scattering over the course of the elution caused by the elution of a buffer component from a previous SEC-SAXS experiment or from the accumulation of materials on the flow cell windows. Regardless, the major component has a profile with a molecular weight estimate that is consistent with an $\alpha_2\beta_2$ tetramer.
